# Noninvasive focal gene transfer of chemogenetic proteins in the primate brain

**DOI:** 10.1101/2025.05.21.655304

**Authors:** M. R. Corigliano, T.V. Parks, S. S. Guretse, L. Letica, I. Zimmermann Rollin, S. K. A. Pell, V. P. Campos, D. Szczupak, D. J. Schaeffer

**Author notes:** Co-first authors (M. R. Corigliano, T.V. Parks). **Corresponding author:** David J. Schaeffer, PhD Assistant Professor Department of Neurobiology University of Pittsburgh Pittsburgh, PA, USA.

## Abstract

The development of chemogenetic neuromodulators, including Designer Receptors Exclusively Activated by Designer Drugs (DREADDs), have enabled focally specific, long-lasting, and reversible neuromodulation in the primate brain. Although systemically delivered synthetic ligands allow for noninvasive actuation of chemogenetic receptors, direct intraparenchymal injection remains atop the available methods to precisely deliver chemogenetic payloads to a specific target of the brain. The requirement of trephination, however, is accompanied by inherent risks of infection, long recovery times, and often tissue damage with concomitant behavioral complications. When considering therapeutic injections, the requirement of transcranial surgery does not translate well to the clinic, especially when repeated administrations are required. Here, we leverage our recent development of transcranial focused ultrasound (tFUS) for noninvasive and focal delivery of adeno-associated viruses (AAVs) carrying excitatory Gq-DREADDs to frontal cortical targets (areas 6DR and 8aD) in the marmoset brain. Using [^18^F]-fluorodeoxyglucose (FDG) positron emission tomography, we demonstrate significant increases in glucose metabolism at the site of viral delivery after administering the DREADD-specific agonist deschloroclozapine (DCZ), as compared to vehicle control. Focal neuronal DREADD expression was confirmed by immunohistochemistry at the site of opening. Through comparison of awake resting-state functional connectivity (whole brain connectivity with the sites of delivery) and structural connectivity (directly injected viral neuronal tracing at the sites of delivery) we demonstrate that the increase in glucose metabolism occurs at both mono- and polysynaptically connected brain regions. Taken together, these results demonstrate the ability to focally deliver excitatory chemogenetics without the need for surgery, allowing for activation of long-range frontal cortex circuits of the primate brain.

## Introduction

Viral-based gene therapies, which leverage the endemic infectivity of a virus to deliver genetic material to cells, hold great promise for the treatment of many otherwise intractable diseases and disorders (see [1] for review). Several viral gene therapies have now received US Food and Drug Administration (FDA) approval [1] with myriad others in development [2]. Although neurological diseases/disorders are being targeted heavily for the development of new gene therapies [2], these treatments will invariably require a method to bypass the blood-brain barrier (BBB) - a selectively restrictive structure regulating molecular access to the central nervous system (CNS) [3, 4]. Generalizable methods facilitating safe penetrance of therapeutics across the BBB, however, remain elusive [3]. Currently, trephination and microinjection are atop the available methods for precise viral vector delivery to the CNS, but present notable health risks including infection and permanent tissue damage [5]. While viral injection does yield robust transduction of neural tissue [6], these potential health complications prevent translation to the clinic at scale [4]. Newly designed adeno-associated virus (AAV) variants possessing the ability to cross the BBB after intravenous (IV) injection [7-13] are an attractive option for delivering genetic material to the CNS, but these constructs are unable to, without additional factors, transduce cells/regions of the brain with specificity - a requirement to avoid mixed or antagonistic effects in potential gene therapies. An emerging new method, transcranial focused ultrasound (tFUS), has been posited as a means to circumvent brain surgery altogether, allowing for the delivery of therapeutics to highly focal brain targets by transiently perturbing the BBB [5, 14-21].

Through the confluence of highly focused sound waves and intravenously injected microbubbles (lipid spheres mechanically agitated by sound), the tight endothelial junctions of the BBB can be transiently opened, allowing for the passage of therapeutics that are otherwise minimally effective at crossing the BBB [4, 22]. Since its inception by Hynynen et al. (2001) [22], tFUS BBB opening has been successfully translated across the phylogenetic spectrum (see the following for demonstrations of tFUS BBB disruption in rabbits [22], pigs [23], mice [24], rats [25], and macaques [20, 26]). We have recently ported the tFUS technique for use in the marmoset monkey (*Callithrix jacchus*), a small New-World primate species that has great promise for the future of neuroscientific modelling of human brain diseases [27]. Despite being roughly the size of a rat (∼350 g as adults), marmosets [21, 28] demonstrate markedly similar functional topologies to those of humans [29-31], and the recent advent of transgenic marmosets [32-34] presents myriad opportunities to test the feasibility of tFUS mediated genetic technologies in nonhuman primate models of disease. To this end, we expanded on our method of BBB perturbation in the marmoset to include the delivery of AAVs carrying fluorescent reporters to marmoset frontal cortex [21] - a concept instantiated via foundational work in rodents [14-18] and recent work in macaques [20]. By incorporating systemic AAV injection into our primate specific tFUS BBB disruption method, we demonstrated focal AAV transduction within insonicated cortex and were able to visualize coursing axons from signal positive neurons [21]. Beyond the delivery of fluorescent reporters, tFUS mediated viral gene delivery has been used in recent reports to reinstate dopamine transmission in a mouse model of Parkinson’s disease [19], suggesting the promise of this technique for the transduction of a variety of translationally relevant neuromodulators. Of these, Designer Receptors Exclusively Activated by Designer Drugs (DREADDs) [35] are promising candidates for noninvasive, reversible circuit modulation.

Modified human muscarinic receptors (DREADDs) have been used extensively in the neurosciences since their introduction in the mid 2000s [35] (see [36] for review). Upon selective activation through administration of their synthetic ligand, these mutant receptors transiently depolarize (Gq) [37] or hyperpolarize (Gi) [38] neurons with concomitant effects on behavior (see [36, 39] for reviews). DREADDs have been deployed to study a variety of processes in the macaque (see [40] for a review of nonhuman primate chemogenetic studies). Indeed, DREADD actuation has been shown to alter macaque working memory [41-45], decision making [44], subjective valuation of reward [46, 47], amygdalar functional connectivity [48], emotional reactivity [49-51], socioemotional attention [50], false belief attribution [52], cutaneous sensation [53], and inhibitory control [54] (see [40] for review). Recently, the efficacy of excitatory DREADDs has also been established in marmoset nonhuman primates via modulation of reward anticipation [55], emotional reactivity [55], and contralateral rotation [56]. Beyond circuit dissection for causal analyses of behavior, the use of DREADDs in nonhuman primates is clinically advantageous as DREADD ligands can be radiolabeled, allowing for *in vivo* assessments of protein expression using positron emission tomography (PET) [41, 43, 45, 47, 49, 51, 53, 56]. Additionally, the extended timescale at which DREADDs are effective allows for noninvasive, circuit-wide indexing of construct function via neuroimaging modalities like [^18^F]-fluorodeoxyglucose (FDG) PET - a method providing quantitative measurements of glucose metabolism as a proxy for neuronal activity [57]. Moreover, DREADD actuators readily cross the BBB [43], reducing the need for repetitive intracranial injections to temporarily excite or inhibit cells of tissues.

Although DREADD ligands can be administered using minimally invasive procedures [41, 43], intracortical injection of viral vectors is currently the preeminent method of focal DREADD delivery to the nonhuman primate brain [40]. Recently, however, tFUS BBB disruption was used to deliver systemically injected AAVs carrying genetic material for either inhibitory or excitatory DREADD receptors to the murine brain [5]. Noninvasively delivered DREADD constructs were demonstrated functional after actuation via increased c-FOS signal (hM3Dq) or inhibition of contextual memory formation (hM4Di) [5]. While tFUS BBB disruption has been shown effective for noninvasive delivery of viral vectors in marmosets [21] and macaques [20], this delivery platform has not yet been used for noninvasive chemogenetic delivery to the primate brain.

Here we demonstrate focal, noninvasive excitatory G(q)-DREADD expression in marmoset frontal cortex via tFUS mediated AAV delivery. Three marmosets received low intensity focused ultrasound to either A8aD (2 marmosets) or A6DR (1 marmoset), transiently perturbing the BBB and allowing for entry of systemically administered AAVs (AAV5, 2 marmosets; AAV9, 1 marmoset) carrying hM3DG(q) to the insonicated neural tissue. After allowing for DREADD expression at the site of BBB opening, we used FDG PET to index changes in glucose metabolism at the site of viral delivery and connected circuitry - a proxy for neuronal activity levels [57] - in the presence of a systemically injected DREADD agonist, deschloroclozapine (DCZ) [43]. As compared to vehicle control, DCZ injections resulted in significant increases in glucose metabolism at the site of DREADD delivery. Immunohistochemical staining revealed robust neuronal DREADD expression localized to the area of BBB perturbation. Through comparison of awake resting-state functional connectivity (whole brain connectivity seeded from the sites of delivery) [58] and structural connectivity (directly injected viral neuronal tracing at the sites of delivery) [59] we demonstrate that the increases in glucose metabolism also occur at both mono- and polysynaptically connected brain regions (to that insonicated) only after DREADD actuation. Taken together, these results demonstrate the ability to focally deliver excitatory chemogenetics – without the need for surgery – allowing for activation of long-range frontal cortex circuits of the primate brain.

## Results

### MRI-based verification of noninvasive BBB Disruption for viral administration

Immediately following a single insonication (aided by microbubble cavitation; see Methods) to either A8aD (Marmosets Pa and V) or A6DR (Marmoset S), each animal was transferred to a 9.4 T MRI and injected with a gadolinium-based contrast agent (GBCA) [58]. The GBCA, in combination with a magnetization-prepared rapid gradient echo (MPRAGE) sequence optimized for the decreased T1 relaxation times in neural tissue exposed to gadolinium, allowed for real-time evaluation of BBB disruption. As demonstrated in our previous work, focal hyperintensities arising from rapid gadolinium extravasation mark the site of insonication [21, 28]. Focal regions of BBB disruption – i.e., GBCA extravasation within the insonication target - were detected in the gray matter of each animal after a single insonication (**Figure 1**). The extent of BBB disruption is consistent with the expected peak of focus of the 1.46 MHz transducer (full width at half maximum of the acoustic pressure distribution) used in these experiments, and between animals for whom the insonication target is similar [21, 28]. For the exact insonication parameters, see the **Methods** below. Immediately after verification of focal BBB disruption (**Figure 1**), each animal was injected (through a catheter placed in the lateral tail vein) with an AAV carrying an excitatory DREADD under the synapsin promoter.

**Figure 1.**
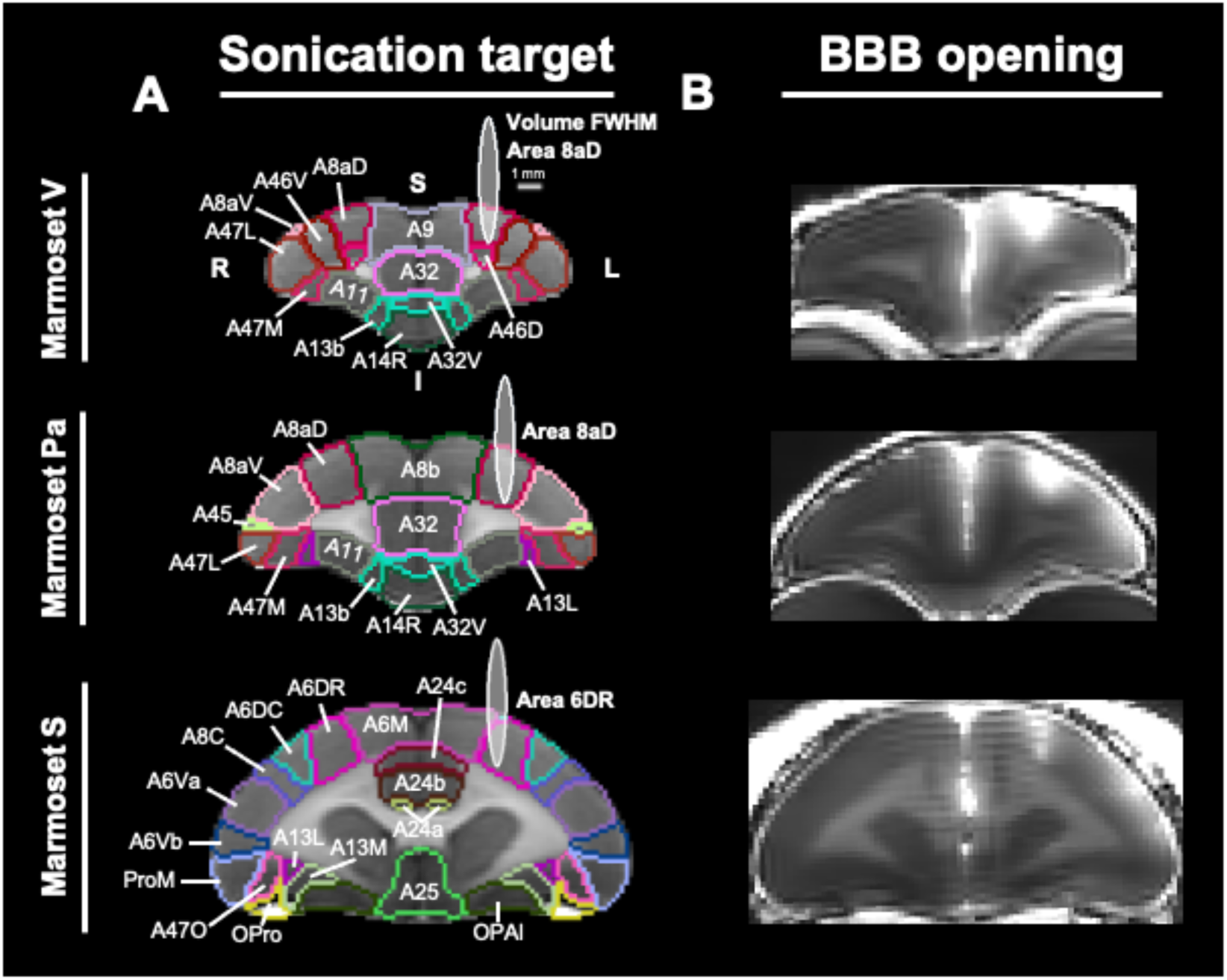
Atlas based targeting of areas 8aD and 6DR for tFUS and the resultant areas of BBB disruption in all marmosets. **(A)** Regions targeted for BBB disruption for all marmosets. Each ellipse (white) represents the theoretical volume of opening for the 1.46 MHz transducer used in this study. Cytoarchitectonic boundaries are based on the Paxinos atlas [62] in Marmoset Brain Mapping template space [58, 61]. **(B)** Regions of BBB disruption for all marmosets visualized via a magnetization prepared - rapid gradient echo (MPRAGE) MR image. The sites of BBB opening are indicated by focal hyperintensities resulting from gadolinium extravasation into the surrounding tissue. Abbreviations: R, Right; L, Left; S, Superior; I, Inferior; A6M, Area 6 of cortex, medial (supplementary motor) part; A6DR, Area 6 of cortex, dorsorostral part; A6DC, Area 6 of cortex, dorsocaudal part; A6Va, Area 6 of cortex, ventral, part a; A6Vb, Area 6 of cortex, ventral, part b; A8b, Area 8b of cortex; A8C, Area 8 of cortex, caudal part; A8aD, Area 8a of cortex, dorsal part; A8aV, Area 8a of cortex, ventral part; A9, Area 9 of cortex; A11, Area 11 of cortex; A13b, Area 13b of cortex; A13L, Area 13 of cortex, lateral part; A13M, Area 13 of cortex, medial part; A14R, Area 14 of cortex, rostral part; A24a, Area 24a of cortex; A24b, Area 24b of cortex; A24c, Area 24c of cortex; A25, Area 25 of cortex; A32, Area 32 of cortex; A32V, Area 32 of cortex, ventral part; A45, Area 45 of cortex; A46D, Area 46 of cortex, dorsal part; A46V, Area 46 of cortex, ventral part; A47L, Area 47 of cortex, lateral part; A47M, Area 47 of cortex, medial part; A47O, Area 47 of cortex, orbital part; ProM, Proisocortical motor region (precentral opercular cortex); OPro, Orbital proisocortex; OPAl, Orbital periallocortex

### Local chemogenetic neuromodulation via noninvasive AAV transduction

Table 1 contains the age of each animal, the viral construct administered after tFUS BBB disruption, and the time allowed for viral construct expression before PET imaging. After allowing for a minimum of two weeks for DREADD expression in the insonicated brain region, marmosets underwent FDG PET imaging either in the presence of DCZ - a potent DREADD agonist with minimal off-target effects on behavior [41, 43] - or a vehicle control (2.5% dimethyl sulfoxide in sterile saline). Compared to vehicle control, we observed significant increases in FDG uptake within the insonicated brain region after systemic injection of DCZ (**Figure 2**) as indicated by Wilcoxon signed rank tests corrected for multiple comparisons (Marmoset V, Insonicated Region adjusted *p* < 0.001; Marmoset Pa, Insonicated Region adjusted *p* < 0.001; Marmoset S, Insonicated Region adjusted *p* < 0.001). Increases in FDG uptake within the insonicated region after DCZ administration were consistent across animals and AAV serotypes used in this study, suggesting the utility of both AAV9 and AAV5 for tFUS mediated gene transfer. Increased FDG uptake was also found in homotopic (non-insonicated) regions of cortex after DCZ administration in all animals (**Figure 2**; Marmoset V, Contralateral Region adjusted *p* < 0.001; Marmoset Pa, Contralateral Region adjusted *p* < 0.001; Marmoset S, Contralateral Region adjusted *p* < 0.001), as detailed in sections below.

**Figure 2.**
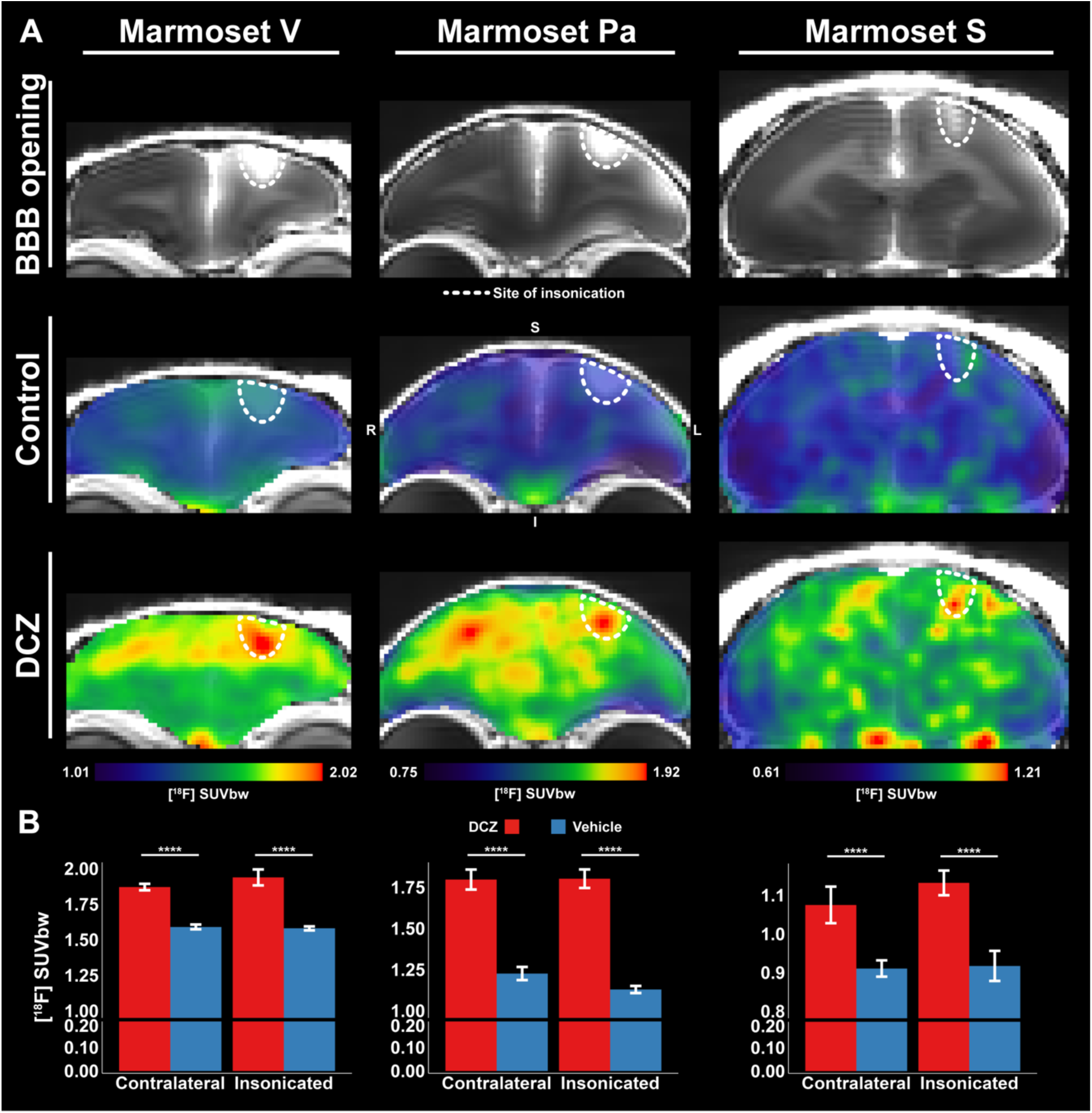
DREADD actuation increases [^18^F]-FDG uptake within the region of BBB disruption and in homotopic cortex. **(A)** Row 1 contains MR images of BBB disruption in each marmoset. White-dotted lines indicate regions of BBB disruption, which are identified through focal hyperintensities resulting from gadolinium extravasation into the surrounding tissue. Rows 2 and 3 display weight-normalized FDG uptake after administration of either a vehicle control solution (row 2) or DCZ (row 3). Warmer colors represent peaks of FDG uptake and are present in both insonicated and homotopic regions after DREADD actuation. **(B)** Bar-graphs showing weight-normalized FDG uptake within both insonicated and homotopic cortex (mean ± standard deviation) after administration of either a vehicle control (blue) or DCZ (red). For all animals, Wilcoxon signed rank tests indicated significant increases in FDG uptake both ipsilateral and contralateral to the insonication site after administration of DCZ as compared to a vehicle control (p < 0.001, ****).

**Table 1.** Subject and viral construct details.

| Marmoset | Age<br>(months) | Weight<br>(g) | Sex | Viral<br>Construct | Titer<br>(GC/mL) | Vol.<br>(μL) | Dose<br>(vg/kg) | Time<br>to<br>Scan<br>(DCZ) |
| --- | --- | --- | --- | --- | --- | --- | --- | --- |
| Pa | 7 | 175 | F | AAV5-SYN1-HA-hM3D(Gq);<br>Addgene<br>121539-AAV5 | $1.1 \times 10^{13}$ | 170 | $1.07 \times 10^{13}$ | 17<br>days |
| V | 51 | 255 | F | AAV9-hSyn-<br>hM3D(Gq)-<br>mCherry;<br>Addgene<br>50474-AAV9 | $2.8 \times 10^{13}$ | 200 | $2.20 \times 10^{13}$ | 27<br>days |
| S | 25 | 260 | F | AAV5-SYN1-HA-hM3D(Gq);<br>Addgene<br>121539-AAV5 | $1.1 \times 10^{13}$ | 170 | $7.19 \times 10^{12}$ | 38<br>days |

### Immunohistochemical validation of DREADD expression

To verify DREADD transduction within the region of BBB disruption, we conducted immunohistochemical analysis on tissue from Marmoset V, who received a construct with the mCherry fluorescent reporter. As seen previously with the tFUS AAV mediated gene transfer method [21], we found robust mCherry expression within the site of insonication (**Figure 3**). Within this 50 µm tissue slice, we observed mCherry fluorescence in 24 cells with clear neuronal morphology covering over 200 µm^2^ of gray matter. Note that mCherry expression was constrained to the site of insonication (**Figure 3**, see **Figure 1** for the extent of BBB disruption in reference to the transducer focus) and not observed in the contralateral hemisphere, which also showed notable increases in FDG uptake after DCZ injection (**Figure 2**).

**Figure 3.**
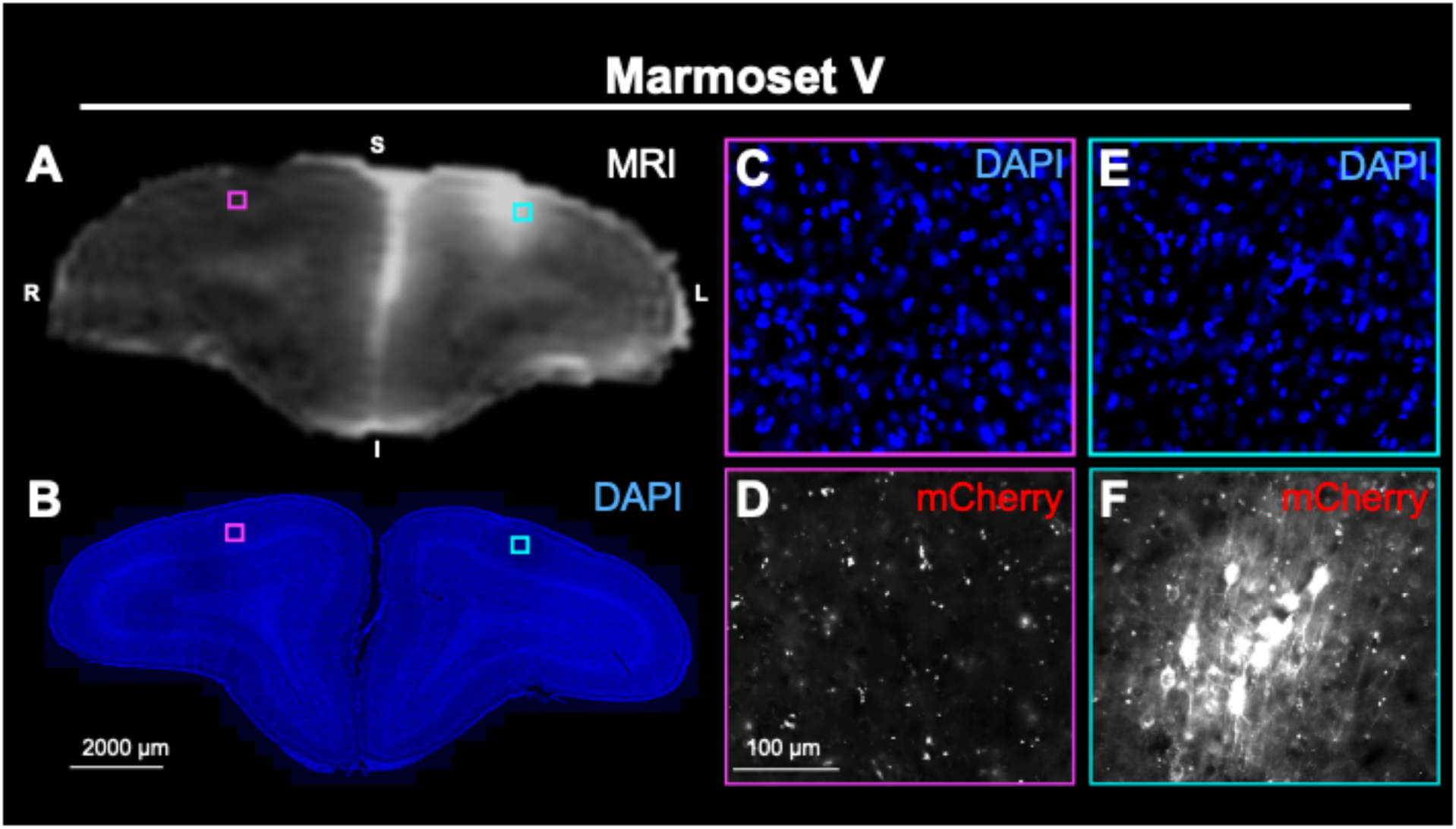
Focal DREADD expression within neurons at the insonicated site. **(A)** MR image of Marmoset V at the region of BBB disruption in area 8aD (left). **(B)** 50 μm tissue section (Marmoset V, 20x magnification) at the coronal level of the MR image above. In both the MR image and coronal tissue slice, cyan and magenta boxes demark the locations of images **(E, F)** and **(C, D)**, respectively. Scale bar = 2000 μm. **(C, D)** Zoomed in view of corresponding tissue region in **(B)** showing positive DAPI signal **(C)** but no mCherry expression **(D)** in cortex contralateral to the site of BBB disruption. Scale bar = 100 μm. **(E, F)** Zoomed in view of corresponding tissue region in image **(B)** showing positive DAPI signal **(E)** and mCherry signal **(F)** within the insonicated cortex. Abbreviations: R, Right; L, Left; S, Superior; I, Inferior

### Comparison of functional and anatomical circuits

Given the increased uptake of FDG in the non-insonicated hemisphere after DREADD activation (**Figure 2**) without concomitant hM3d(Gq) expression in this same locus (see **Figure 3D**), we examined population functional [58] and individual long-range structural connections [59] of insonicated regions via open-source marmoset resources. As an index of structural connectivity (specifically, monosynaptic anterograde connections), data from anterograde neuronal tracer injections included in The Brain/MINDS Marmoset Connectivity Resource [59] were chosen for analysis based on their proximity to the insonication targets presented in this study (Marmoset V, A8aD, injection R01-0054; Marmoset Pa, A8aD, injection R01-0059; Marmoset S, A6DR, injection R04-0079) [59]. Voxel-wise functional connectivity profiles of each insonication point were downloaded from our Marmoset Brain Connectome resource, a resting-state functional magnetic resonance imaging (RSfMRI) resource generated with data from 32 marmosets [58]. To demonstrate DCZ-related activation at homotopic sites of connection, **Figure 4** compares these (injection [59] and RSfMRI [58]) data to FDG uptake after DREADD activation in each animal in a coronal plane at their site of insonication. Given the focal transduction of AAVs within the insonicated site (see **Figure 3** and previous work [21]), we expected areas of increased radiotracer uptake outside of the insonication target to be functionally connected, anatomically connected, or both functionally and anatomically connected to the target itself. Indeed, **Figure 4** demonstrates consistent overlap between foci of increased glucose utilization, functional connectivity peaks, and anterograde neuronal projections for each insonication location. Overlap between FDG peaks, functional connectivity peaks and anterograde projection targets is most apparent in regions homotopic to those insonicated but is also found in other frontal areas including 8b and A6M (**Figure 4**).

**Figure 4.**
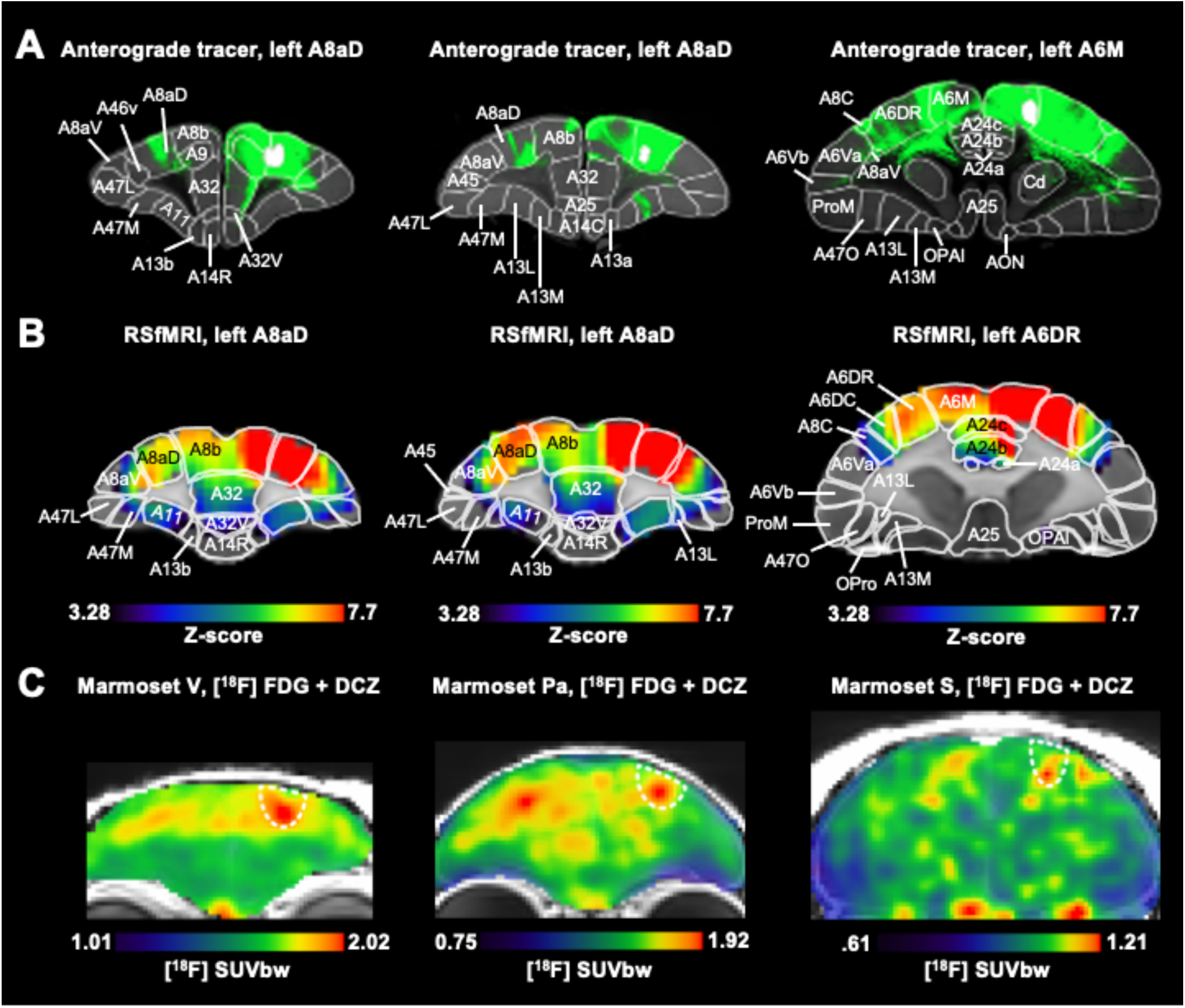
Comparison of anterograde tracing, RSfMRI, and [^18^F]-FDG PET data for each insonicated region. **(A)** Anterograde tracer data collected from The Brain/MINDS Marmoset Connectivity Resource [59] demonstrating structural connectivity from injections closest to insonication targets (Marmoset V, injection R01-0054; Marmoset Pa, injection R01-0059; Marmoset S, injection R04-0079. **(B)** Resting-state functional connectivity maps collected from the Marmoset Brain Connectome (Z-score of 3.28 corresponds to p < 0.001) [58]. Single voxel seeds were chosen based on the location BBB disruption. Labels in rows 1 and 2 are derived from the Paxinos atlas parcellation [62]. **(C)** Weight-normalized FDG uptake after DCZ injection for all animals. White dotted lines indicate the extent of BBB disruption in the coronal plane. Abbreviations: A6M, Area 6 of cortex, medial (supplementary motor) part; A6DR, Area 6 of cortex, dorsorostral part; A6DC, Area 6 of cortex, dorsocaudal part; A6Va, Area 6 of cortex, ventral, part a; A6Vb, Area 6 of cortex, ventral, part b; A8b, Area 8b of cortex; A8C, Area 8 of cortex, caudal part; A8aD, Area 8a of cortex, dorsal part; A8aV, Area 8a of cortex, ventral part; A9, Area 9 of cortex; A11, Area 11 of cortex; A13a, Area 13a of cortex; A13b, Area 13b of cortex; A13L, Area 13 of cortex, lateral part; A13M, Area 13 of cortex, medial part; A14C, Area 14 of cortex, caudal part; A14R, Area 14 of cortex, rostral part; A24a, Area 24a of cortex; A24b, Area 24b of cortex; A24c, Area 24c of cortex; A25, Area 25 of cortex; A32, Area 32 of cortex; A32V, Area 32 of cortex, ventral part; A45, Area 45 of cortex; A46D, Area 46 of cortex, dorsal part; A46V, Area 46 of cortex, ventral part; A47L, Area 47 of cortex, lateral part; A47M, Area 47 of cortex, medial part; A47O, Area 47 of cortex, orbital part; AON, Anterior olfactory nucleus; Cd, Caudate nucleus; ProM, Proisocortical motor region (precentral opercular cortex); OPro, Orbital proisocortex; OPAl, Orbital periallocortex

### Long-range chemogenetic neuromodulation via noninvasive AAV transduction

Because we observed increased activation - measured via FDG uptake - in homologous regions in the contralateral hemisphere after DREADD activation, we hypothesized that we would observe similar increases in activation within other brain areas functionally connected to the insonication target. To test this, we employed the ‘fingerprinting’ technique, which has been used previously to compare patterns of activity across a circuity (i.e., interareal connectivity) [31], rather than comparing intensity values with any single region. By vectorizing and scaling the patterns (here patterns of FDG uptake across connected regions), the fingerprint plots allow for visual comparison of overall patterns of activity. Using resting state functional connectivity maps of insonicated regions collected from the Marmoset Brain Connectome [58], we placed regions of interest (ROIs) along nodes of the functional circuit (as identified by sites of peak seed-based correlation values) of each insonication location (**Figure 5**). From these ROIs, we extracted weight-normalized FDG uptake from functionally connected cortical/subcortical locations for each animal/condition to generate condition-dependent activation fingerprints. As expected, FDG uptake in regions connected to the insonicated location was elevated after DCZ administration as compared to vehicle injections (**Figure 5**). Importantly, in marmosets Pa and S, changes in FDG uptake between DCZ and vehicle controls were lowest for the entorhinal cortex, a control region neither functionally (Z-score < 2) nor directly structurally connected (lack of fluorescent reporter) to insonication targets in A8aD and A6DR. Indeed, elevations of activity in DCZ trials were apparent in nearly every functionally connected region to that receiving BBB disruption, especially areas receiving dense projections from insonicated targets [60]. Albeit at a lower magnitude than the connected sites, ROIs containing no monosynaptic input from insonicated targets - as evidenced by minimal coincidence between RSfMRI/anterograde projection maps within ROIs themselves (DICE score < 0.2) - also showed increases in FDG uptake after DCZ administration (**Figure 5**). In ROIs receiving minimal anterograde input (excluding entorhinal ‘control’ areas), resting state functional connectivity remained robust and, like the functional connectivity of monosynaptic connections, qualitatively similar to PET fingerprints (**Figure 5**), suggesting large-scale interhemispheric increases in activation after chemogenetic modulation of a single target region.

**Figure 5.**
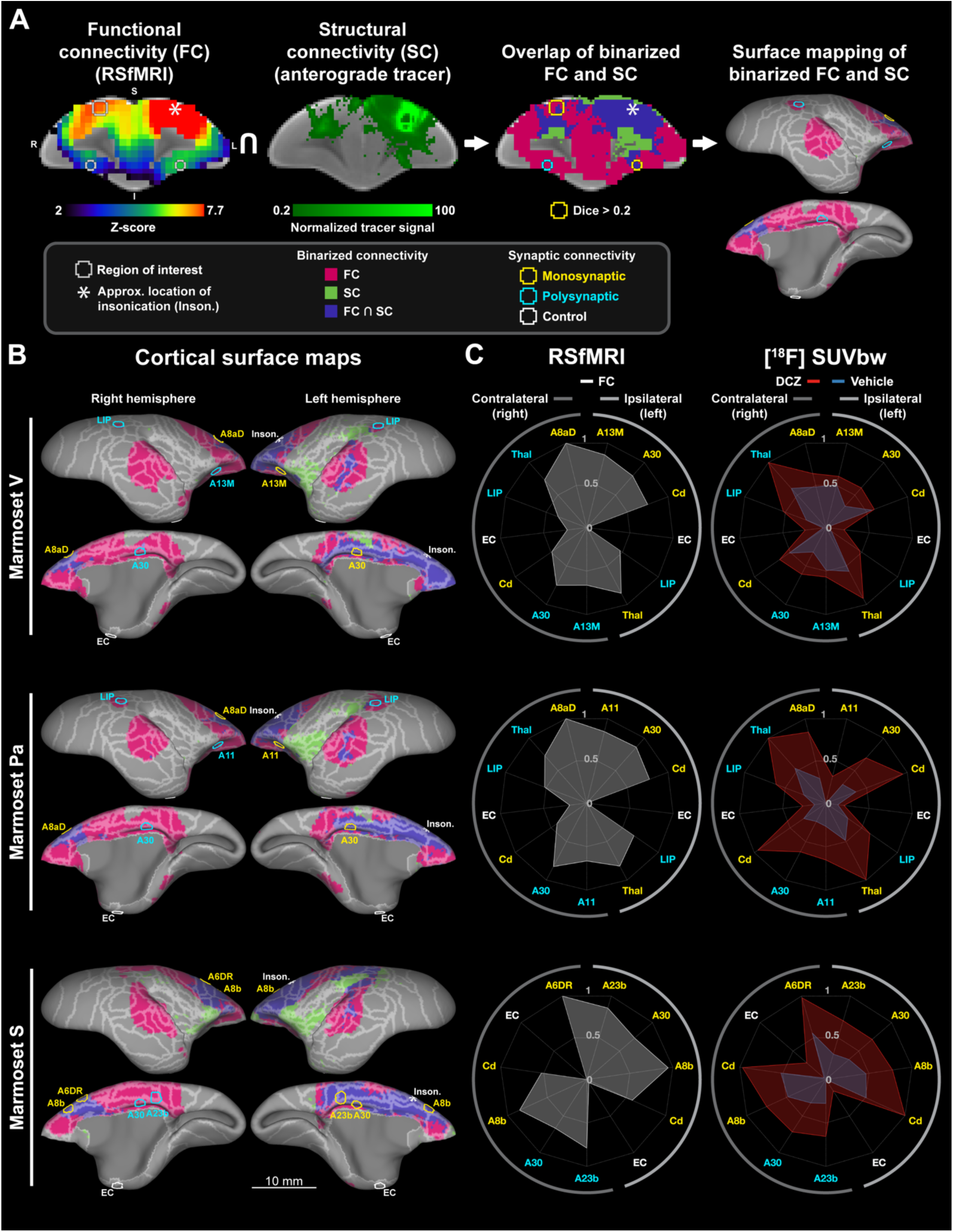
DCZ-mediated increases in glucose uptake are found across regions functionally connected to insonicated cortex independent of the presence of monosynaptic anterograde projections. **(A)** Schematic outlining the process of categorizing the structural connectivity between areas of BBB disruption and functionally connected ROIs. The leftmost image displays resting-state functional connectivity with the insonicated region (white asterisk) along with 1.5 mm diameter spherical ROIs (gray circles) centered at connectivity peaks (and mirrored to the contralateral hemisphere). RSfMRI [58] and viral anterograde tracer data (from injections near to insonication targets; image 2) [59] were binarized and combined to create overlap volumes (image 3) - e.g. a visual representation of regions exclusively functionally connected (magenta) OR structurally connected (green) as well as regions functionally AND structurally connected (purple) with an insonication target. Within each ROI, binarized functional and structural data were used to calculate Sørensen–Dice coefficient similarity (Dice score > 0.2 = monosynaptic anterograde connectivity (yellow circles); Dice score < 0.2 = polysynaptic connectivity (cyan circles)). An example overlap volume (image 4) and categorized ROIs on the surface of the marmoset brain [61]. **(B)** Surface maps of functional (magenta), structural (green) or both functional and structural (purple) connectivity with the region of BBB disruption (white asterisk) for each animal. White borders on the surface of the brain are from the Paxinos atlas [62] in Marmoset Brain Mapping space [61]. ROIs - represented as outlines - are also included in surface maps for each animal., The color of ROIs corresponds to their designation as monosynaptic (yellow), polysynaptic (cyan) or as a control region (white) demonstrating no marked structural or functional connectivity with the insonicated region. Scale bar = 10 mm. **(C)** RSfMRI (left) and PET (right) interareal fingerprints for each animal included in this study. Functional connectivity (white outline) and weight-normalized FDG (vehicle control = blue outline; DCZ = red outline) data are independently scaled between 0 (lowest values) and 1 (highest values) to facilitate comparisons of patterns between modalities (RSfMRI vs PET) and experimental trials (DCZ vs vehicle control). As in the surface maps, the nature of structural connectivity between each ROI and its corresponding insonication target is reflected in the color of its label (see above). ROI location relative to the area of BBB disruption is denoted by enclosing semi-circles (dark gray = contralateral hemisphere, light gray = ipsilateral hemisphere). Abbreviations: A6DR, Area 6 of cortex, dorsorostral part; A8aD, Area 8a of cortex, dorsal part; A8b, Area 8b of cortex; A11, Area 11 of cortex; A13M, Area 13 of cortex, medial part; A23b, Area 23b of cortex; A30, Area 30 of cortex; Cd, Caudate nucleus; Thal, Thalamus; LIP, Lateral intraparietal area of cortex; EC, Entorhinal Cortex

## Methods

### Animals

Data were collected from three adult female marmosets (*Callithrix jacchus*; see Table 1 for subject details) housed at the University of Pittsburgh. Female animals were chosen for these foundational experiments simply on availability for terminal procedures. Marmosets were either opposite-sex pair housed or family group housed, fed a diet consisting of twice daily provisions of commercial chow including purified diet, and supplemented with fresh fruit and vegetables daily with drinking water provided ad libitum. Foraging materials and enrichment were provided daily. Prior to tFUS, MRI and PET procedures, marmosets received intramuscular injections of meloxicam (0.2 mg/kg; 07-893-5916, Patterson Vet Supply, Inc., Loveland, CO, USA) and glycopyrrolate (0.005 mg/kg; 1226607, McKesson Medical-Surgical, Richmond, VA, USA). Animals were anesthetized (induced and maintained) with 2% isoflurane delivered via mask and fitted with a 26-guage venous catheter for tFUS, MRI, and PET procedures. During the procedures, heart rate, blood oxygen, respiration, and rectal temperature were monitored. Body temperature was maintained with heated water blankets or heated air. All experimental procedures were in accordance with ethical guidelines for animal testing established by the University of Pittsburgh Institutional Animal Care and Use Committee.

### Focused Ultrasound BBB Disruption

To transiently disrupt the BBB in a focal region of cortex for noninvasive viral vector delivery, marmosets were placed in an RK-50 Marmoset tFUS apparatus (FUS Instruments Incorporated, Toronto, ON, Canada) incorporating a marmoset stereotax (Model SR-AC; Narishige International Incorporated, Amityville, New York, USA) and an MRI-based based atlas for stereotactic targeting (https://www.marmosetbrainconnectome.org [58, 61, 62]). The tFUS apparatus used here is described in detail in [28] and summarized in [21]. Briefly, an automated 3-axis position system guided a 55 mm spherically focused 1.46 MHz transducer (FUS Instruments Incorporated, Toronto, ON, Canada) that was rigidly mounted to the positioning system to allow for precise control of the sonication location. The number of pulses, repetition period, and number of bursts commanded by the software were generated by an external waveform generator (Siglent SDG 1032X, Siglent Technologies, Solon, Ohio, USA) and sent through a 15 W amplifier to the transducer. The transducer was sealed using a 3D printed cover incorporating an o-ring seal and a polyimide film face that held degassed water. The transducer and cover were immersed in a ∼300 mL tank containing degassed water, the polyimide bottom of which was coupled to marmoset head using degassed, warmed ultrasound transmission gel (01-50, Parker Laboratories, INC., Fairfield, NJ, USA). Water degassing was accomplished using a FUS-DS-50 portable water degasser (FUS Instruments Incorporated, Toronto, ON, Canada).

Transcranial insonication was based on stereotactic position of which the zero point (x, y, z = 0, 0, 0) corresponded to a location equidistant from the center of the ear bars and in the axial plane of the inferior most aspect of the orbit bars. tFUS positioning software (Morpheus framework; FUS Instruments Incorporated, Toronto, ON, Canada), which incorporates the marmosetbrainconnectome.org [58] and the marmosetbrainmapping.org [63] atlases, was used to select predefined insonication targets based on the Paxinos parcellation of the marmoset brain [62]. These locations were further refined based on preprocedural MRI (using the same parameters defined below). Two regions were selected for sonication, area 8aD for its role in controlling eye movements in the marmoset [64, 65], and area 6DR which, in addition to its role in the neural control of eye movements [64, 65], is a central functional connectivity hub in the marmoset [66, 67].

At the beginning of all tFUS procedures, the marmoset scalp was cleared of hair using a combination of electric clippers and depilatory cream (Nair Shower Cream, Church & Dwight Co, Inc., Ewing, NJ, USA). Immediately prior to insonication, a bolus of lipid microbubbles (350 µL/kg; Definity, Lantheus Medical Imaging, Billerica, MA, USA) - prepared by mixing 0.86 mL of sterile saline with 0.1 mL of microbubbles - was administered via the implanted catheter to aide in BBB disruption. To ensure complete clearance of the microbubble solution from the catheter hub, a flush of 200 µL sterile saline was administered to all marmosets. Following microbubble injection, the 1.46 MHz transducer was oriented directly above the insonication target for a single sonication with the following parameters: acoustic pressure = 1.27 - 1.38 MPa (estimated with 47% deration [28]), burst duration = 20-25 ms, burst period = 1000 ms, number of bursts = 60 bursts.

### MRI Acquisition and Denoising

To visualize BBB opening after insonication, anesthetized marmosets were transferred from the tFUS apparatus to a 9.4 T 30 cm horizontal bore MRI scanner (Bruker BioSpin Corp, Billerica, MA, USA), equipped with a Bruker BioSpec Avance Neo console and the software package Paravision-360 (version 3.6; Bruker BioSpin Corp, Billerica, MA, USA). Marmosets were placed into a custom-designed MRI-compatible stereotax for imaging (see https://www.marmosetbrainconnectome.org/apparatus/ [68] for computer-aided design files), after which 100 μL of a gadolinium-based MRI contrast agent (GBCA; Gadavist^TM^, gadobutrol; Bayer Healthcare Pharmaceuticals, Leverkusen, Germany) was systemically injected through the implanted catheter. GBCA was cleared from the catheter hub using 200 µL of sterile saline. During imaging, radiofrequency transmission was accomplished with a custom 135 mm inner diameter coil. A custom-built 14-channel phased-array marmoset-specific coil was used for radiofrequency receiving [68]. A magnetization prepared - rapid gradient echo (MPRAGE) sequence was used to detect reduced T1 relaxation times resulting from gadolinium extravasation at the site of tFUS BBB permeation with the following parameters: TR = 6,000 ms, TE = 3.29 ms, field of view = 42 x 35 x 25 mm, matrix size = 168 x 140 x 100, voxel size = 0.250 x 0.250 x 0.250 mm, bandwidth = 50 kHz, flip angle = 12 degrees, total scan time = 20 minutes, 6 seconds. To enhance the signal-to-noise ratio (SNR) of the anatomical MR images, we applied a denoising procedure as in [69, 70]. We used a variance-stabilizing transformation (VST) framework tailored for Rician-distributed noise, characteristic of magnitude MR images [71]. The forward VST was applied to transform the heteroscedastic Rician noise into approximately homoscedastic Gaussian noise. The transformed images were then denoised using the block-matching and 4D filtering (BM4D) algorithm [72]. Finally, the inverse VST was applied to restore the denoised images to their original intensity range.

### Systemic AAV injections

Immediately after BBB opening was verified by GBCA extravasation at the targeted site, either AAV9 or AAV5 (Addgene, Watertown, MA, USA; see Table 1 for details and dosages) were injected as a single bolus through the implanted catheter. Following AAV injection, a 200 µL flush of sterile saline cleared virus from the catheter hub. At the conclusion of experiments, the scalp was covered in Aquaphor ointment (Beiersdorf Inc., Wilton, CT, USA) to prevent irritation from hair removal. After animals sufficiently recovered from anesthesia, they were returned to their home cages.

### [^18^F]-FDG PET Imaging of DREADD Actuation

In the weeks following viral delivery (Marmoset Pa, 17 days; Marmoset V, 27 days; Marmoset S, 38 days), all marmosets underwent anesthetized FDG PET scans to noninvasively probe the effects of DREADD actuation. Marmosets were fixed in stereotactic plane using a custom-built marmoset PET stereotax (https://www.marmosetbrainconnectome.org/apparatus/ [68]) and fit with a 26-gauge venous catheter. Immediately prior to FDG imaging, animals received either a single dose of DCZ (100 µg/kg) or 2.5% dimethyl sulfoxide (DMSO, D2650-100ML, Sigma-Aldrich, Inc., St. Louis, MO, USA) diluted in sterile saline (vehicle solution) via the implanted catheter. Two preparations of DCZ were used in this study: 1) DCZ powder (SML3651-5MG, Sigma-Aldrich, Inc., St. Louis, MO, USA) dissolved in DMSO at a concentration of 2 mg/mL, and then diluted to a final concentration of 2.5% DMSO in sterile saline (Marmosets Pa and S) or 2) DCZ dihydrochloride powder (HB9126-10mg, Hello Bio Inc, Princeton, NJ, USA) dissolved in distilled water at a concentration of 1 mg/mL (Marmoset V). At the time of DCZ/vehicle solution administration, marmosets also received a bolus of FDG at a dose of 80 MBq (2.16 mCi). Immediately following both radio-and-synthetic ligand dosing, PET data were collected using a Bruker Si-78 small animal PET/CT (Bruker BioSpin Corp, Billerica, MA, USA), with a console running ParaVision 360 (version 3.6, Bruker BioSpin Corp, Billerica, MA, USA). PET scans were acquired over 90-minutes with a field-of-view of 120 x 80 x 80 mm. PET scans were dynamically reconstructed using a maximum *a posteriori* projection in the following bins: 3x20; 4x30; 2x60; 5x300; 6x600 seconds. At the conclusion of PET imaging, a computed tomography (CT) image was acquired to aide in image registration with the following parameters: field of view = 79.6 x 81.3 mm, pixel size = 200 µm, X-ray source filter = 1mm aluminum, frame averages = 1, scanning mode = step and shoot, rotation angle = 1°.

### [^18^F]-FDG PET Image Processing and Analysis

To prepare data for an analysis of FDG uptake after DREADD actuation, all PET and MRI scans were preprocessed using PMOD quantification software (version 4.4, PMOD Technologies LLC, Zurich, Switzerland). In brief, the anatomical orientation of each marmoset in their PET images was verified using lateralized soft and bony tissue landmarks (PMOD’s View). The orientation of the brain in anatomical MR images was verified using the site of insonication - a feature demarcating the left hemisphere (PMOD’s View). PET scans were time corrected (PMOD’s View) and brought into alignment with their anatomical MR image (PMOD’s Fusion). After initial alignment, PET images from each animal were rigidly registered to their anatomical MRI (PMOD’s Fusion). PET scans were then averaged 30 minutes to 90 minutes (bins 15 to 20, PMOD’s Fusion) and converted to standard uptake value normalized to body weight (SUVbw) images (PMOD’s Fusion). At the conclusion of pre-processing, PET images were converted from digital imaging in communications and medicine (DICOM) format to neuroimaging informatics technology initiative (NIfTI) files using the PMOD View tool. Post-tFUS MRI images were converted to NIfTI format in Analysis of Functional NeuroImages (AFNI, version AFNI_23.0.03) software (dcm2niix_afni) [73-75] for further analysis outside of PMOD.

Comparisons of FDG uptake within insonicated and contralateral (homotopic) region between DCZ and vehicle trials were conducted with pre-processed PET scans in native space. Using PMOD software (PMOD’s Fusion), 1.5 mm diameter spheres were placed in the region of BBB disruption and mirrored to the contralateral (non-insonicated) region of cortex. Values from each pixel within insonicated and contralateral non-insonicated ROIs were extracted (PMOD’s Fusion, Pixel Dump), pre-processed, and visualized via plot of mean and standard deviation using R (RStudio, 2023.06.2+561, Posit Software, PBC) [76-81]. In R, data were assessed for normality via univariate Shapiro-Wilk tests (shapiro_test [82]) with *p* < 0.05 indicating a non-normal distribution. Due to a high prevalence of non-normal distributions (data not shown), comparisons of weight-normalized FDG uptake within insonicated and homotopic (non-insonicated) regions between trials (DCZ and vehicle) were conducted using a paired samples Wilcoxon signed rank test in R (wilcox_test) [82]; a *p* < 0.05 - adjusted for multiple comparisons using the Holm method - indicated a significant difference between trial type. Explicitly, statistical tests used pixel values from pre-processed scans (one scan per condition) in native space and were calculated across voxels, not calculated across animals or insonication locations.

### RSfMRI and [^18^F]-FDG PET Fingerprinting Analysis

To examine the effects of DREADD actuation within the functional circuit of each insonication target, we generated region specific fingerprints [31, 83-85] using open access RSfMRI data [58] and FDG PET data collected for this manuscript. Preprocessed, MPRAGE images were skull-stripped using UNet Studio software (version for Windows 7+) [86] and linearly registered to a T1-weighted template image [58, 63] using the FMRIB Software Library’s (FSL) FLIRT command [87, 88]. MR images in template space were then nonlinearly registered to the same template [58, 63] using Advanced Normalization Tools (ANTs) [89]. Transformation matrices from both linear and nonlinear registration were saved for each animal and applied to their respective PET images yielding files for group analysis.

ROIs were selected using RSfMRI correlation maps generated by selection of a single voxel within each animal’s insonicated region via the Marmoset Brain Connectome webpage (https://www.marmosetbrainconnectome.org) - an open access resource of RSfMRI data from 32 marmosets [58]. Regions were selected based on the presence of robust functional connectivity peaks (i.e., high blood oxygen level dependent time course correlation values) with the insonicated region (threshold = Z-score > 2). A control region was also selected for each animal based on a lack of both significant correlation and anticorrelation with the insonicated region (threshold = within Z-score -2 to 2). Note that each point was mirrored in the contralateral hemisphere after selection yielding ROIs in both the right and left hemisphere of the brain. Based on these criteria, 11-13 ROIs were selected for each animal included in this study. At each point, we generated a 1.5 mm diameter sphere (FSL’s fslmaths) from which an average value was extracted (AFNI’s 3dmaskave) from RSfMRI and PET images resampled to the T1-weighted template image referenced above (AFNI’s 3dresample). Average RSfMRI and PET (DCZ and vehicle control trials) values were rescaled from 0 (lowest functional connectivity or FDG uptake) to 1 (highest functional connectivity or FDG uptake) in Matlab (version 9.14.0.2206163 (R2023a), MathWorks, Natick, MA, USA). Rescaling data allowed us to compare patterns of activation between modalities directly when presented in a radar plot (a customized version of radarplot [90]).

### Determination of Structural Connectivity Type

To examine the anatomical underpinnings of insonicated region functional circuits, we compared the topographies of RSfMRI correlation maps [58] and viral anterograde tracer data [59] downloaded from open access web-based resources. Anterograde tracing data were collected from three injections included in The Brain/MINDS Marmoset Connectivity Resource [59]. Injection locations were chosen based on their proximity to the insonication target of each marmoset; two injections from A8aD (Marmosets Pa and V) and one from A6M (approximating insonication of A6DR of Marmoset S). Injection selections, by marmoset, were as follows: Marmoset V, thy1-tTA 1/TRE-clover 1/TRE-Vamp2mPFC, 0.2 µl injected, injection R01-0054; Marmoset Pa, thy1-tTA 1/TRE-clover 1/TRE-Vamp2mPFC, 0.2 µl injected, injection R01-0059; Marmoset S, thy1-tTA 1/TRE-clover 1/TRE-Vamp2mPFC, 0.2 µl injected, injection R04-0079. See https://dataportal.brainminds.jp/marmoset-tracer-injection/viewer [59] for further injection details. Tracer data were then registered to the Marmoset Brain Mapping template [58, 63] using ANTs (antsApplyTransforms). RSfMRI maps collected for each insonication location (see ***RSfMRI and [^18^F]-FDG PET Fingerprinting Analysis***) were then resampled to the tracer data in template space (AFNI’s 3dresample). Thresholds of 3.28 (Z-score, *p* < 0.001 for RSfMRI maps [58]) and 0.2 were applied to functional and tracer data, respectively, to reduce noise. After applying these thresholds, functional and anatomical connectivity data were binarized (AFNI’s 3dcalc). Sørensen–Dice indices (Dice score) were calculated between binarized structural and functional connectivity maps within ROIs defined for fingerprinting analyses (AFNI’s 3ddot). ROIs were defined as monosynaptic anterograde connections if Dice scores were greater than 0.2. To visualize the location of ROIs in relation to resting-state functional and anterograde structural connectivity, overlap maps were generated in Connectome Workbench (version 1.5.0) [91] and, along with ROIs, displayed on surface maps of the marmoset brain [61].

### Immunofluorescence, Microscopy and Cell Counting

To conduct immunofluorescence staining, Marmoset V was euthanized via injection of a solution containing 390 mg/mL pentobarbital sodium and 50 mg/mL phenytoin sodium (100 mg/kg, EUTHASOL, Virbac, Westlake, TX, USA). Then we perfused transcardially with cold phosphate buffer saline (PBS) followed by 4% paraformaldehyde (PFA). The brain was extracted, post-fixed in 4% PFA, and cryoprotected in ascending concentrations of sucrose (10% - 30%). Following cryoprotection, the brain was embedded in tissue freezing medium (TFM-C, General Data Healthcare, Cincinnati, OH, USA), frozen, and sectioned at 50 µm via a Leica CM1950 cryostat (Leica Biosystems, Deer Park, IL, USA). Sections were stored at -20 °C in cryoprotectant solution consisting of 30% glycerol, 30% etholene glycol, 30% distilled water and 10% pH 7.2 0.2M phosphate buffer. To begin immunofluorescence staining, cryoprotectant solution was removed from free floating sections via a series of 1x PBS washes (3 x 10 minutes). Sections were then gently rocked in blocking solution (5% Donkey Serum + 1% Bovine Serum Albumin + 0.2% Triton-X + PBS) at room temperature for 60 minutes. Following the blocking step, sections were incubated at 4 °C in primary antibody solution containing a rat anti-mCherry amplifier conjugated to a 594 nm fluorophore (1:250, M11240, ThermoFisher Scientific, Waltham, MA, USA), mouse anti-NeuN (1:500, MAB377, MilliporeSigma, Burlington, MA, USA), 1% Donkey Serum and 1x PBS. Sections were gently agitated in primary antibody solution for a total of 72 hours. Upon completion of the primary antibody incubation, tissue sections were washed in 1x PBS (3 x 10 minutes) and gently agitated in secondary antibody solution containing Alexa Fluor^TM^ donkey anti-mouse 488 (1:1000, A21202, ThermoFisher Scientific, Waltham, MA, USA), 1% donkey serum and 1x PBS for 1 hour at room temperature. All tissue sections were incubated with DAPI (4’,6-Diamidino-2-Phenylindole, 1:1000, 62248, ThermoFisher Scientific, Waltham, MA, USA) in 1x PBS for 10 minutes followed by additional 1x PBS washes (2 x 10 minutes). Tissue sections were mounted on Superfrost™ Plus slides (22037246, ThermoFisher Scientific, Waltham, MA, USA) and allowed to dry. Slides were coverslipped using Immu-Mount (9990402, ThermoFisher Scientific, Waltham, MA, USA) prior to imaging.

To quantify the extent of transduction resulting from focal BBB disruption, stained slices were imaged on an AxioImager M2 epifluorescence microscope (Carl Zeiss, White Plains, NY, USA). Acquisition settings were kept constant for all images within a staining group. Margins for tiled layout images were hand delineated. Large-scale tiled images were obtained at 20x and stitched using ZEN microscopy software (ZEN 3.5, blue edition, Zeiss, Oberkochen, Germany). As described above, insonicated regions were located *in vivo* via extravasation of GBCA into the surrounding neuropil. An MR image demonstrating the locus of BBB disruption in Marmoset V was used to identify this same location in sliced tissue. In the region of BBB opening, cells with clear neuronal morphology that were also positive for mCherry were manually counted.

## Discussion

Here we demonstrate noninvasive, focal transduction of functional excitatory chemogenetic proteins in marmoset frontal cortex. Using tFUS aided by systemically injected microbubbles, we transiently and focally disrupted the BBB, allowing for passage of viral constructs that transduced neurons with excitatory G(q)-DREADD proteins (**Figure 1**, see [21, 28] for our previous work establishing this technique in the marmoset). Excitatory DREADDs were activated by noninvasively administered ligands and, using radiolabeled glucose as a proxy for neural activity [57], we demonstrated modulation of frontal cortex regions in three nonhuman primates (**Figure 2**). Results from FDG PET experiments were supported by immunohistochemistry showing selective reporter expression at the site of insonication (**Figure 3**). Increased glucose metabolism was also observed in contralateral homotopic connections (**Figure 4**) as well as other connected circuitry (**Figure 5**) after DCZ administration. Using fully awake RSfMRI [58] and direct intracortical injection data from anterograde tracers [59], we demonstrate that DCZ-driven increases in glucose metabolism were present at both mono- and polysynaptically connected regions as compared to unconnected (control) regions (**Figure 5**). Taken together, these data demonstrate the efficacy of tFUS BBB disruption for noninvasive delivery of functional neuromodulators that, when actuated, alter activation levels across the functional circuit of insonication targets – including functionally connected regions not receiving monosynaptic projections from regions in which the BBB was disrupted.

The central focus of this study was to expand tFUS mediated viral gene delivery to include neuromodulators with both scientific and clinical applications; moreover, to demonstrate that tFUS mediated gene delivery and modulation is viable in frontal cortex targets, with other extant reports (apart from our demonstration of FUS AAV delivery in marmosets [21]) targeting subcortical regions [20]. To this end, three marmosets received a single cortical insonication resulting in focal BBB disruption (**Figure 1**). Immediately following verification of BBB opening with GBCA MRI, marmosets were systemically injected with either AAV5 or AAV9 constructed to express the excitatory DREADD receptors: hM3d(Gq). After allowing several weeks for viral expression, we conducted FDG PET on the three marmosets in two conditions: administration of the DREADD agonist DCZ [43] or a vehicle control. As compared to vehicle injection trials, FDG uptake was significantly higher within insonicated regions after systemic DCZ administration (**Figures 2**). These results confirm those of Mimura and colleagues (2021) [56], who demonstrated increased relative FDG uptake at the target of a unilateral DREADD injection following receptor actuation.

In support of our finding of increased FDG uptake within insonicated regions, immunohistochemical staining of tissue from Marmoset V revealed neuronal cell bodies positive for mCherry that were limited to the region of BBB disruption (**Figure 3**). These data indicate successful noninvasive DREADD protein expression within a focal region of the primate brain. While recent works have advanced the use of tFUS BBB disruption to facilitate AAV based gene transfer, these studies generated openings either broad in spatial extent and/or within rodents [5, 14-20]. Here, we demonstrate BBB disruption and transduction patterns with precision on the order of millimeters. Although restricted in spatial extent, we do not anticipate the DREADD expression profile presented here to limit the potential utility of tFUS mediated viral gene delivery. Reliable alterations in macaque behavior have been shown after minimal cellular DREADD expression [46, 92]. Consequently, we consider the spatially precise pattern of DREADD transduction resulting from tFUS BBB opening to be advantageous for precise manipulation of neural circuits. Even so, the effects observed in this study may be further improved via experimentation with new AAV serotypes, especially those specifically engineered for use with tFUS BBB disruption and detargeted from peripheral organs [93] - a preferable property due to the promiscuity of many AAV serotypes [94, 95]. Work with enhancer elements, such as those designed to limit AAV expression to cortical inhibitory interneurons [96], may also be advantageous to broaden the potential impact of this technique for both circuit modulation and the treatment of a variety of diseases/disorders (see [97] for review).

In addition to increased FDG uptake at the site of insonication, we also observed marked elevations in FDG uptake within homotopic regions of cortex (relative to those insonicated) after DCZ injection (**Figures 2).** To explore these results further, we examined the functional and structural connectivity of insonicated targets. Using The Brain/MINDS Marmoset Connectivity Resource [59], the Marmoset Brain Connectome [58], and PET data presented here, we compared the topography of anterograde projections from injection regions approximating insonicated cortex, group resting-state functional connectivity profiles from single voxels within insonicated regions, and individual animal FDG uptake data in the coronal plane (**Figure 4**). Overlap between anterograde structural connectivity, resting state functional connectivity, and DCZ-mediated elevations in FDG uptake were high for all animals (**Figure 4**); in combination with the focal cellular expression of hMD3(Gq) only within insonicated cortex (**Figure 3**), these data suggest monosynaptic communication between homotopic regions may be responsible for the propagation of increased activation levels after DCZ administration. These results are in line with observations from our lab of transgene expression in anterograde projections after tFUS mediated viral gene delivery [21]. Albeit mediated by a different viral construct, these results are also consistent with those seen during an optogenetic neuromodulation paradigm whereby stimulation of cell bodies in one hemisphere could elicit increases in oxidative metabolism within homotopic cortex [98].

The multimodal combination of open access RSfMRI data [58] and tracer data [59] also enabled us to assess whether DCZ mediated alterations in glucose uptake were present in other regions functionally connected with insonication targets. By comparing overlap between functional and structural connectivity, we deduced whether ROIs received monosynaptic anterograde projections from areas in which the BBB was perturbed. Explicitly, we used a measure of overlap (Dice score) between functional and structural connectivity maps from each insonication location to determine if functionally connected regions (to the site of BBB disruption) also received monosynaptic anterograde projections from insonication targets (Dice score > 0.2 indicated a monosynaptic anterograde projection, Dice score < 0.2 indicated a polysynaptic connection; see ***Determination of Structural Connectivity Type***). In contrast to regions contralateral to the site of BBB disruption, regions ipsilateral to the insonication point presented high correspondence between anterograde projections and peaks of functional connectivity - i.e. were monosynaptically connected with the region of insonication (**Figure 5**). DCZ mediated increases of glucose uptake were apparent in nearly all regions receiving monosynaptic projections from insonicated targets (**Figure 5**). These data support the findings of Michaelides and colleagues (2013) [57], who demonstrated that local DREADD activation can modulate FDG uptake in connected regions of the rat brain. Further, we observed increases in FDG uptake after DREADD actuation in ROIs where overlap between functional and anatomical connectivity was low, evidencing modulation of both structural (monosynaptic anterograde) and functional circuits after insonication of only a single target region (**Figure 5**) - findings consistent with other nonhuman primate studies (see paragraph below). While our findings of both structural and functional circuit modulation rely on our assumptions used to identify monosynaptic projection targets, the nature of the open access tracer data we employed [59] does not allow us to make definitive claims on the possibility that regions are monosynaptically connected in a retrograde fashion. It is also critical to note that ROIs were selected using group resting state connectivity maps [58], while tracer data from each injection stem from a single animal [59] precluding our ability to account for inter-animal variations in structural connectivity [99]. Indeed, inter-animal variability may also account for points where results we present here diverge from regions canonically viewed to be monosynaptically connected (**Figure 5**) - e.g. A8aD and the lateral intraparietal area (LIP) [60, 100] - as peaks in group functional connectivity maps may not reflect the exact point of entry of projections from cells captured in a single injection.

As is evident in **Figure 2**, FDG uptake increased most dramatically at the site of viral delivery and in connected circuity, but there were also more subtle widespread increases (comparing vehicle and DCZ scans for each animal, **Figure 2**). Using fMRI, others have noted altered functional connectivity between regions structurally connected and structurally unconnected to injection targets after DREADD actuation [48, 53]. These results are complemented by those of Hirabayashi and colleagues (2024) [45] who, using functional PET, found that chemogenetic inhibition of the orbitofrontal cortex modulated task related activation in other structurally/functionally connected regions of the macaque brain. Further, Hirabayashi et al. (2024) [45] demonstrate that DREADD-mediated reductions in activity of orbitofrontal cortex are sufficient to abolish delay related activity for preferred stimuli in anterior ventral temporal cortex - an anatomically connected region - during an object memory task. Although we utilize an excitatory G(q)-DREADD construct in our experiments, we note a qualitatively similar spread of modulated activity from the site of insonication, particularly in connected regions (**Figures 2, 4** and **5**). Indeed, the prospect of spreading depolarization in response to actuation of focal excitatory chemogenetics may be of great importance to adequately place in context any causal findings from behavioral paradigms incorporating DREADDs. Nevertheless, similarities between results in macaques and those presented here are particularly notable given the difference in method employed by each study - e.g. tFUS mediated viral gene delivery - a noninvasive technique - recapitulates effects observed after direct intracranial injections.

While we were unable to test for DREADD mediated behavioral alterations in this study as all animals were selected based on a pre-designation for humane euthanasia (not allowing sufficient time for behavioral training), we specifically selected insonication targets based on their behavioral and translational relevance. Indeed, both insonicated regions presented in this study are involved in the control of saccades in marmosets [64, 65]. Future studies validating robust and reliable behavioral effects will be critical for demonstrating the validity of the approach described here. Behavioral validation notwithstanding, our results demonstrate the feasibility of an approach that is dramatically simpler than performing a neurosurgical injection and that can be completed in less than two minutes. Moreover, this report is one of the first to demonstrate noninvasive, focal AAV delivery to primate cortex and we look forward to a formal assessment on how tFUS mediated viral gene delivery can lead to the eventual modulation of well characterized behaviors.

In this work, we present the use of tFUS and lipid microspheres to disrupt discrete regions of the BBB in marmoset frontal cortex, promoting focal, excitatory DREADD transduction via systemically delivered viral constructs. Across three marmosets, activation of Gq-DREADD constructs via IV DCZ administration consistently increased radiolabeled glucose metabolism not only within insonicated regions, but in mono- and polysynaptically (as informed by resting-state functional connectivity and anterograde viral tracing) connected circuitry both ipsilateral and contralateral to the site of BBB disruption as well. Taken together, these results reinforce the feasibility of tFUS BBB opening for the delivery of biologics in marmosets [21] and represent a completely noninvasive suite of tools for chemogenetic circuit modulation in nonhuman primates. For these reasons, we expect this technique can revolutionize the field of nonhuman primate chemogenetics by increasing the feasibility with which other labs can access this foundational method of circuit modulation. Moreover, this technique may empower the design of brain-region based, noninvasive, gene therapies for the treatment of complex neurological diseases and disorders.

## Acknowledgements

We wish to thank Brianne L. Stein, Lauren Dubberley, and Dr. Julia Oluoch for animal care and preparation. This work was supported by the National Institute of Neurological Disorders and Stroke of the National Institutes of Health under award number R21NS125372 (D.J.S.).

